# Methodological choices strongly modulate the sensitivity and specificity of lesion-symptom mapping analyses

**DOI:** 10.1101/2025.11.04.686650

**Authors:** Margaret Jane Moore, Chris Rorden, Gail A. Robinson, Jason B. Mattingley, Nele Demeyere

**Affiliations:** Queensland Brain Institute, University of Queensland, St. Lucia, Queensland; Department of Psychology, University of South Carolina, Colombia, SC, USA; School of Psychology, University of Queensland, Brisbane, Australia; Canadian Institute for Advanced Research, Toronto, Canada; Nuffield Department of Clinical Neurosciences, University of Oxford, Oxford, United Kingdom

**Keywords:** Lesion-symptom Mapping, Multivariate Lesion Mapping, Stroke, Multiverse Analysis, Neuroimaging

## Abstract

Lesion mapping results can vary substantially as a function of specific analysis parameters, but the extent to which individual methodological choices interact to modulate the sensitivity and specificity of results is not clear. Here, we employed a large-scale simulation approach to inform practical recommendations for lesion symptom mapping studies.

Routine clinical imaging from 959 stroke survivors (mean age = 72.5, 49.3% female) was used to conduct 384,780 lesion mapping analyses based on simulated behavioural data. Each simulated analysis used different combinations of plausible sample inclusion criteria, analysis parameters (e.g., correction factors), analysis types (e.g., univariate vs. multivariate), and underlying target correlates. Simulated analysis accuracy (Dice similarity coefficient and percent coverage of target correlates) was compared across designs.

Overall, analysis accuracy varied widely and was substantially modulated by the specific design used. Analyses that maximised lesion coverage by including large and diverse samples reliably outperformed analyses using more restricted samples. Analyses using direct total lesion volume controls outperformed analyses using other (or no) volume corrections across all accuracy measures. False discovery rate corrections yielded the best performance in terms of target coverage, while permutation corrections yielded the best Dice coefficients. While multivariate approaches were more accurate than univariate analyses in terms of Dice coefficient, univariate analyses generated higher target hit rates and percent target coverage.

These results identify specific analysis designs suitable for studies aiming to maximise their sensitivity and/or specificity to underlying critical correlates, while highlighting the inferential strengths and weaknesses of these complementary approaches.

## Introduction

Lesion-symptom mapping is a popular analytical approach in which the location of brain damage is used to infer causal brain-behaviour relationships (Bates et al., 2003; de Haan & Karnath, 2018; Moore, Demeyere, et al., 2023). This method identifies brain regions are associated with impairments when injured and associated with intact function when spared (Bates et al., 2003; de Haan & Karnath, 2018; Moore, Demeyere, et al., 2023; Rorden & Karnath, 2004). Lesion-symptom mapping methods have been implemented to localise the neural correlates of a wide range of fundamental cognitive functions (Moore et al., 2024; Weaver et al., 2021). However, recent research has provided evidence that the accuracy of unconstrained lesion-symptom mapping analyses may be spatially biased (Inoue et al., 2014; Mah et al., 2014), and that results can vary substantially as a function of specific analysis choices (Ivanova et al., 2021; Mirman et al., 2018; Sperber et al., 2019). The aim of the current study was to use a large-scale simulation approach to extend understanding of how specific analytical choices interact to modulate the outcomes of lesion symptom mapping analyses.

Lesion symptom mapping, like any other statistical approach, entails an inherent trade-off between sensitivity and specificity (Kimberg et al., 2007; Sperber, 2022). Lesion symptom mapping analyses aiming to optimize sensitivity may employ more liberal statistical approaches to prioritize the detection of significant neural correlates over minimizing the risk of voxel-wise false positives (Sperber, 2022). Analyses aiming to maximise specificity may opt for more conservative methods to identify “core” correlates over minimizing the risk of false negative results (Moore et al., 2024; Moore & Demeyere, 2022). Alternatively, researchers may wish to take a balanced statistical approach. Each of these methodological choices are well-suited to addressing specific types of research questions in lesion symptom mapping analyses, but the specific methodological choices required for each of these statistical approaches remains unclear.

When planning any lesion symptom mapping analysis, researchers must select from many design options which each have the potential to modulate the sensitivity and specificity of the resultant analysis. For example, researchers need to choose sample inclusion/exclusion criteria by considering (or not considering) factors such as the type and location of the lesion, imaging modality, and lesion numerosity (de Haan & Karnath, 2018; Moore, Demeyere, et al., 2023). Each of these choices has potentially critical consequences. For example, the spatial distribution (and degree of overlap) of lesions within study samples ultimately constrains the range of brain areas analyses are able to draw conclusions about (Moore et al., 2023). As sample selectivity increases, the range of neural areas which can be tested with sufficient statistical power decreases (Kimberg et al., 2007). Researchers may therefore aim to maximise lesion coverage by using lenient exclusion criteria (e.g., including patients with multiple spatially distinct lesion sites or different imaging modalities), but it is unclear whether the added noise associated with these choices outweighs the benefit of including a larger sample.

Researchers must also select between an extensive range of potential statistical approaches (e.g. univariate versus multivariate), covariate controls, multiple comparison corrections, and minimum lesion overlap inclusion criteria (Ivanova et al., 2021; Mirman et al., 2018; Sperber & Karnath, 2017). Past research has highlighted several key analysis choices that influence outcomes, but it is not yet clear what the optimal analysis choices are. For example, analyses which include statistical corrections for lesion volume generally yield improved specificity relative to analyses which do not, but this gain occurs at the cost of sensitivity (DeMarco & Turkeltaub, 2018). Moreover, the magnitude of this improvement differs across different volume correction types, with some types of volume corrections argued to be too conservative (DeMarco & Turkeltaub, 2018; Zhang et al., 2014). Applying lesion volume corrections results in higher specificity which may be in line with the goals of some, but not all, study designs (Moore et al., 2024; Sperber, 2022). There is similar uncertainty surrounding the optimal method for correcting for multiple comparisons. While False Discovery Rate (FDR) corrections are commonly used in lesion symptom mapping (Mirman et al., 2018), these corrections generate very liberal results which may be spatially displaced from true, underlying neural correlates, especially in cases where lesion volume is not controlled for (Mirman et al., 2018). Studies aiming to prioritize specificity generally opt for either family-wise error rate via permutation or Bonferroni corrections, but these approaches may be too conservative (de Haan & Karnath, 2018; Mirman et al., 2018). There is a similar lack of consensus surrounding voxel-level minimum inclusion criteria, with different studies arguing the advantages and disadvantages of restricting analysis to voxels impacted by sufficient lesions (e.g. 10% of the total sample) (Sperber & Karnath, 2017).

While traditional lesion symptom mapping approaches generally use mass-univariate statistical approaches, multivariate approaches are rising in popularity (DeMarco & Turkeltaub, 2018; Moore, Demeyere, et al., 2023; Pustina et al., 2018; Zhang et al., 2014). Mass-univariate approaches consider each voxel independently and are therefore theoretically unable to distinguish between critical neural correlates and non-critical areas that are likely to be damaged by the same lesions (Mah et al., 2014). Multivariate lesion symptom mapping approaches address this limitation by considering all voxels concurrently to generate a single, robust prediction of impairment (DeMarco & Turkeltaub, 2018; Mah et al., 2014; Moore, Demeyere, et al., 2023; Pustina et al., 2018). However, it is not yet clear whether multivariate approaches reliably outperform univariate approaches in terms of sensitivity and specificity.

Previous work has addressed this issue by examining the extent to which the significant voxels yielded by lesion mapping analyses agree with a pre-defined, ground-truth target area using a range of accuracy measures such as dice similarity coefficient, percentage of target voxels reported as significant, and distance between significant results and target areas (Ivanova et al., 2021; Mah et al., 2014; Pustina et al., 2018). While some studies have reported improved accuracy in multivariate analyses (e.g., Pustina et al., 2018), several large-scale simulation studies have reported no clear accuracy difference between multivariate and univariate approaches (Ivanova et al., 2021; Sperber et al., 2019). Additionally, the accuracy of multivariate approaches appears to be highly variable when different analysis parameters (e.g. lesion volume) and sample sizes are used (DeMarco & Turkeltaub, 2018; Sperber et al., 2019). The accuracy of multivariate analyses is also often quantified in terms of impairment prediction accuracy, rather than the extent to which resulting significant voxels align with underlying “true” neural correlates (Griffis et al., 2024). However, robust impairment predictions can be supported by voxels that are not underlying critical impairments (Griffis et al., 2023; Moore et al., 2023). For example, the presence of unilateral damage to right hemisphere voxels could act as an informative predictor of the absence of impairment on a picture naming task (e.g. nominal aphasia). This relationship could theoretically lead a multivariate model to identify the degree of damage to right hemisphere areas as the strongest predictor of picture naming impairment, in the absence of a causal relationship between damage to these areas and this deficit. For this reason, it remains unclear what the optimal statistical approach for lesion symptom mapping is.

In addition to explicit analytical choices, past simulation studies have shown that sample size and lesion coverage characteristics also strongly modulate analysis accuracy. The accuracy of simulated lesion symptom mapping has generally been found to improve as the sample size (i.e. number of considered lesions) and degree of overlap at key regions of interest increases (Ivanova et al., 2021; Moore et al., 2023). Past studies have aimed to maximise sample sizes by combining different neuroimaging modalities and including comparatively low-resolution routine clinical imaging (e.g. CT). However, the benefits and drawbacks of this approach relative to study designs that employ exclusively higher resolution neuroimaging (e.g. MRI) have not yet been systematically evaluated (de Haan & Karnath, 2018). Lesion symptom mapping results are ultimately determined by the spatial topography of the included lesion overlay and underlying critical correlates, but the extent to which this association varies across different samples (and sample sizes) remains unclear (Mah et al., 2014).

Simulation-based lesion symptom mapping approaches can provide unique insights into the impact of design choices on analysis accuracy (Ivanova et al., 2021; Mah et al., 2014; Moore et al., 2022; Sperber et al., 2019). Specifically, a large number of potential datasets and corresponding analysis designs can be generated by making each possible permutation of reasonable methodological choices (Dragicevic et al., 2019; Ivanova et al., 2021). Each of these potential experimental designs can then be simulated, allowing for cumulative results to be evaluated to identify factors leading to results variability. In the context of lesion symptom mapping, simulation approaches enable investigation of many different analysis types, analysis parameters, and sample factors that may ultimately modulate analysis accuracy. Here we employed a simulation-based approach to address this key knowledge gap and to support practical recommendations for lesion symptom mapping studies aiming to optimize outcomes. The findings provide a highly detailed insight into how the choices made in lesion symptom mapping designs ultimately impact accuracy and offer guidance for future lesion symptom mapping studies.

## Methods

### Stroke Neuroimaging Data

We report a retrospective analysis of existing routine stroke neuroimaging data collected as a component of the Oxford Cognitive Screening Program (NHS REC reference 14/LO/0648, 18/SC/0550, 12/WM/00335) and by The University of Queensland’s Neuropsychology Clinic (Metro South and Metro North Queensland Health and the University of Queensland HREC/16/QPAH/793). Participants were included in the present study if they had clinical imaging depicting visible stroke lesions, and that the scan was of sufficient quality to support lesion delineation and spatial normalisation. All participants provided informed written consent (in line with the Declaration of Helsinki). Data from 959 stroke patients (mean age = 72.5 (SD = 13.3), 49.3% female, 8.0% left-handed) was included in the present study.

Lesion masks were generated in line with the standard processing protocol reported by (Moore, 2022). Specifically, lesions were manually delineated on native space axial scan slices by a trained expert (MJM) (765 CT, 194 MR (135 T2, 1 T1, 58 FLAIR). This imaging was collected a mean of 14.4 days post stroke (SD = 68.2, range = 0-924). Stroke types were reported as 61.4% Ischemic, 33.3% Haemorrhagic, and 5.3% unreported, and lesion sides were reported as 37.8% left, 50.2% right, 18.3% bilateral.

Masks were smoothed at 5mm full width at half maximum in the z-direction and binarised using MRIcron (Rorden, 2007). Scans and lesions were then reoriented to the anterior commissure to reduce transformation degrees of freedom and warped into 1×1x1 mm MNI space using nonlinear normalisation and age-matched templates (SPM, Ashburner et al., 2016; Clinical Toolbox, Rorden et al., 2012). Normalised scans and lesions were visually inspected for accuracy prior to inclusion. All scans that could not be accurately normalised accurately were excluded.

### Simulation analysis

The voxels underlying real-world neuropsychological deficits are ultimately unknown, meaning that simulation-based lesion symptom mapping approaches are required to measure analysis accuracy (Ivanova et al., 2021; Mah et al., 2014; Moore et al., 2022). In this approach, simulated behavioural scores are generated from real lesion data by defining the extent to which each considered lesion overlaps with a pre-defined critical target site (e.g. target ROIs). These critical sites include regions that are commonly impacted in vascular-related neurological injuries, as well as regions that are less commonly damaged in stroke. These simulated behavioural scores can then be input into lesion symptom mapping analyses, allowing for accuracy to be quantified by comparing the resultant significant voxels to the known target site.

This approach was used in two separate simulation analyses (univariate and multivariate) to investigate how different analysis parameters modulate results accuracy. Target ROIs were generated by creating an array of spheres (volume = 4.2 cm^3^) covering the brain (Figure 1, B). This volume and shape were arbitrarily selected to ensure adequate coverage while balancing the computational resources needed to evaluate performance at each target. Real patient lesion masks were compared with each target to simulate impairment severities (the percent of target ROI voxels impacted by each lesion). Targets impacted by fewer than 3 lesions were not included in analyses, resulting in 671 target areas.

**Figure 1:**
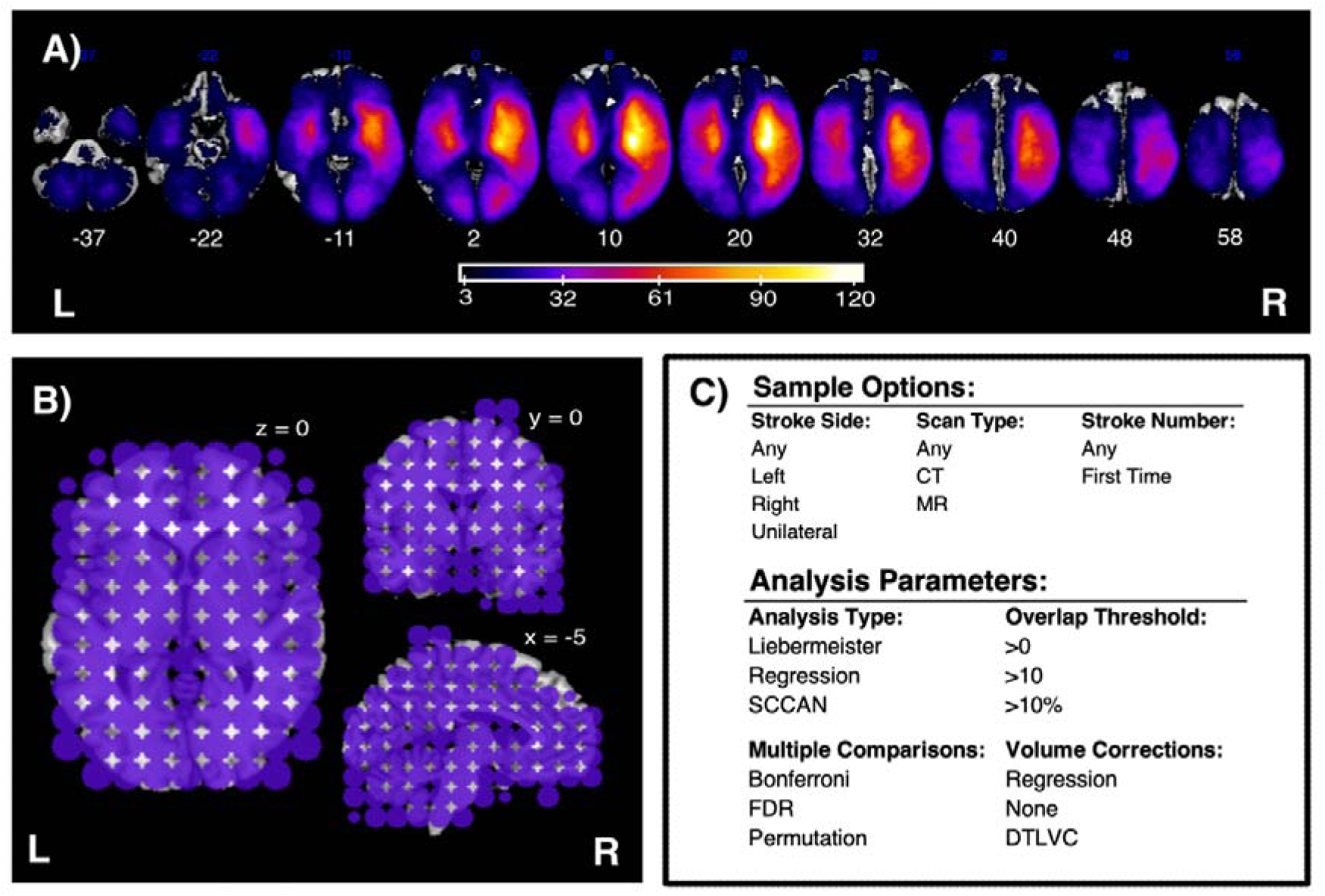
A visualisation of the lesion data and parameters used in this study’s simulation analyses. (A) Lesion overlay plot for the full sample (n = 959). Colour denotes number of overlapping lesions at each location. Only areas impacted in at least 3 lesions are visualised. MNI slices −37 – 58 are presented. (B) The anatomical targets used in this study. Only targets impacted in at least three lesions are included in this visualisation. (C) All patient sample and analysis parameter options considered in the univariate analyses. The specific parameters considered in multivariate simulations are reported in text.

Simulation analysis accuracy was quantified using two metrics: percent target coverage and Dice coefficient. Percent target coverage is the proportion of target ROI voxels which are significant in simulated analysis. Percent coverage prioritises capturing sensitivity (i.e. detecting the underlying correlate) over specificity, as this metric does not penalise false positives or overestimation of target volume. Conversely, Dice coefficient provides a stricter measurement of the agreement between two binary image segmentations. Unlike percent coverage, Dice penalises voxel masks for both over- and under-estimating the underlying target area. For this reason, Dice captures both voxel-wise sensitivity and specificity. Percent coverage and Dice have a U-shaped relationship, such that low Dice coefficients occur both when percent coverage is very high (prone to overestimation) and very low (underestimation) (See Supplementary Figure 1).

All simulation analyses were implemented using LESYMAP (Pustina, 2019) in R and were run using The University of Queensland’s High Performance Computing Cluster (Dell EMC Technologies Australia Pty Ltd).

### Univariate Simulation Analyses

The purpose of the univariate simulation analyses was to investigate how different sample inclusion criteria and analysis parameters modulate the accuracy of lesion symptom mapping analyses. Within these analyses, behavioural scores were simulated using the approach described above but the patients included in each analysis (sample inclusion criteria) and the lesion symptom mapping approach used (analysis parameters) was systematically varied across simulated analyses.

Possible sample inclusion criteria included all possible combinations of stroke side, scan type, and stroke history inclusion criteria (Figure 1, C). Within stroke side, *Left/Right* categories include all patients with damage confined to the relevant brain hemisphere, *Unilateral* includes patients with lesions confined to a single hemisphere, and *unselected* include all patients, regardless of lesion location. Within scan type, *CT/MR* include patients with the relevant imaging type while *Unselected* include all patients regardless of imaging modality. Within stroke history, *First Stroke* includes only patients exhibiting evidence of a single stroke event while *Unselected* analyses include all patients regardless of stroke numerosity.

Analysis parameter options were generated by varying analysis type, multiple comparison corrections, minimum overlap threshold, and lesion volume corrections. Within univariate simulation analysis types, Chi Squared (with Yates correction) and regression analyses were used. Chi Squared analyses test binary associations between voxel impairment (spared/impaired) and behavioural scores (spared/impaired). Within these analyses, lesions which overlapped with the relevant behavioural target were scored as impaired while all other patients were considered unimpaired. Chi-squared analyses were included as the binary analysis option in this study as other common approaches (e.g. Liebermeister tests) are not implemented in LESYMAP (Pustina, 2019; Rorden et al., 2009). In cases where voxel values are binary (spared/impaired), the regression analyses used are equivalent to non-parametric t-tests but have the added benefit of allowing voxel-level corrections for multiple comparisons (described below) (Pustina, 2019). In cases where these corrections are used, the regression approach correlates continuous behavioural scores with volume-corrected lesion values (Pustina, 2019).

Three different multiple comparison corrections (Bonferroni, False Discovery Rate (FDR), and family-wise error rate permutation) and 3 minimum overlap thresholds (voxels impacted in at least one patient (*>0*), voxels impacted in at least 10 patients (*>10*), and voxels impacted in at least *10%* of the sample) were considered. Within lesion volume corrections, *None* analyses employed no correction for lesion volume, *Regression* analyses included volume as a covariate of no interest in analysis, and *DTLVC* analyses employed direct total lesion volume control (DTLVC, Zhang et al., 2014) corrections which weight voxel values by lesion volume. When impossible analysis parameters combinations were excluded (e.g., Chi-squared analyses with volume corrections), these analysis parameters yielded 33 unique analysis designs. Each of these potential analysis designs was applied for each target ROI (n=759) and sample groups (n = 24) yielding 601,128 unique potential univariate simulation analyses.

The results generated from these simulated lesion symptom mapping analyses were then cumulatively analysed to measure the extent to which analysis accuracy (quantified in terms of percent target hits, Dice, and percent coverage) varied across different analysis designs. Specifically, accuracy was compared across underlying anatomical targets. Multivariate regression analyses were conducted to determine the extent to which general analysis factors (e.g. sample size, average lesion volume, number of lesions impacting the target, results cluster size) modulated accuracy. Analysis accuracy was then compared across individual sample inclusion and analysis parameter choices using ANOVA tests (Bonferroni corrected for multiple comparisons). Next, multivariate regression analyses were conducted to evaluate whether individual analysis designs modulated accuracy when general analysis factors were considered. In these analyses, the use of each individual analysis parameter (binarised) was added to a multivariate regression predicting results accuracy from general analysis factors. In cases where added analysis choices significantly and positively explained accuracy variance when considered alongside general analysis factors, these analysis choices were considered to be associated with improved accuracy. Throughout the Results section, all reported R^2^ values are adjusted.

### Multivariate Simulation Analyses

Next, multivariate simulation analyses were conducted to allow for the accuracy of multivariate and univariate lesion symptom mapping approaches to be compared. Specifically, the accuracy of simulated sparce canonical correlation-based lesion symptom mapping (SCCAN) was compared with that of matched univariate simulations (described above). SCCAN is a popular multivariate lesion symptom mapping approach that builds a best-fit model leveraging multivariate lesion patterns to predict behavioural scores. The voxels retained in this model represent the predicted neural correlates associated with behavioural scores (Pustina et al., 2018). SCCAN uses a modified L1 penalty to handle highly correlated input variables (Pustina et al., 2018). The percentage of input predictors included in final SCCAN solutions was optimised through an iterative sparseness optimisation and cross-validation procedure (Pustina et al., 2018). Using this approach, SCCAN models are considered significant if a model’s predicted behavioural scores are above chance (p < 0.05) within cross-validation.

Notably, this sparseness optimisation procedure is computationally intensive, meaning that it is not feasible to simulate all possible analyses. This issue increases exponentially as the number of voxels (and lesion masks) included in the analysis increases (Pustina et al., 2018). This issue was managed by reducing the number of factors considered in multivariate simulations (relative to univariate simulations). First, to reduce the maximum sample size and number of voxels considered, only patients with unilateral stroke damage (i.e., exclusively left or right hemisphere damage) were included in multivariate simulations, yielding two possible sample groups. All other sample inclusion criteria (e.g. scan type, stroke number) were not considered in multivariate simulations. In terms of analysis parameters, potential minimum overlap thresholds were reduced to >10 and 10% to further minimize the number of voxels considered in analysis. Volume correction options were reduced to *regression* and *no corrections*, as DLVTC corrections cannot be implemented in SCCAN analyses (Pustina, 2019). SCCAN does not employ corrections for multiple comparisons, and voxel-wise significance is dependent on model fit (Pustina et al., 2018). These factors yielded 8 possible analysis designs which were applied over the 759 behavioural targets, resulting in 6072 potential simulated multivariate lesion symptom mapping analyses. The data were used to evaluate the relative accuracy of multivariate approaches by comparing the results of these simulations with those of univariate approaches using matched analysis parameters and samples.

### Pre-Registration and Data Availability

This study’s simulation analysis design and core analyses were pre-registered on the Open Science Framework (https://osf.io/6kcha/registrations). All code and simulated data associated with the project are openly available on the Open Science Framework (https://osf.io/6kcha/). Lesion masks are available on request from the authors.

## LSM Multi Results

### Univariate Analyses General Descriptives

63.4% of the potential univariate analysis designs met criteria for inclusion in the simulation analysis. Of these analyses, 72.0% yielded significant results, 66.7% of which overlapped with the underlying target. The average significant voxel cluster size was 11.5 cm^3^ (sd = 16.1, 0.001-130.4). Within analyses yielding significant results, the average Dice was 0.05 (sd = 0.08, range =0 – 0.73) and the average target coverage was 43.6% (sd = 41.6, range = 0-100).

Analysis accuracy varied across the target ROIs (Figure 2). Of the analyses producing significant results, the probability of hitting the underlying target roughly aligned with the degree of statistical power present in the lesion distribution. Hit probability was highest within MCA and PCA stroke territory and lowest in regions less commonly impacted in the sample (e.g., the most anterior frontal regions, peripheral cerebellar regions, and regions spanning the midline) (Figure 2, B). This accuracy distribution was mirrored in the average target coverage, with very high coverage in areas with high lesion overlap and poor accuracy in less affected areas (Figure 2, C). Results Dice was highest within the pons, bilateral basal ganglia, and cerebellum (Figure 2, C). Each of these regions is commonly impacted by small, relatively spatially homogeneous lesions (e.g., basal ganglia lacunar strokes) (Bassetti et al., 1996; Wardlaw, 2005).

**Figure 2:**
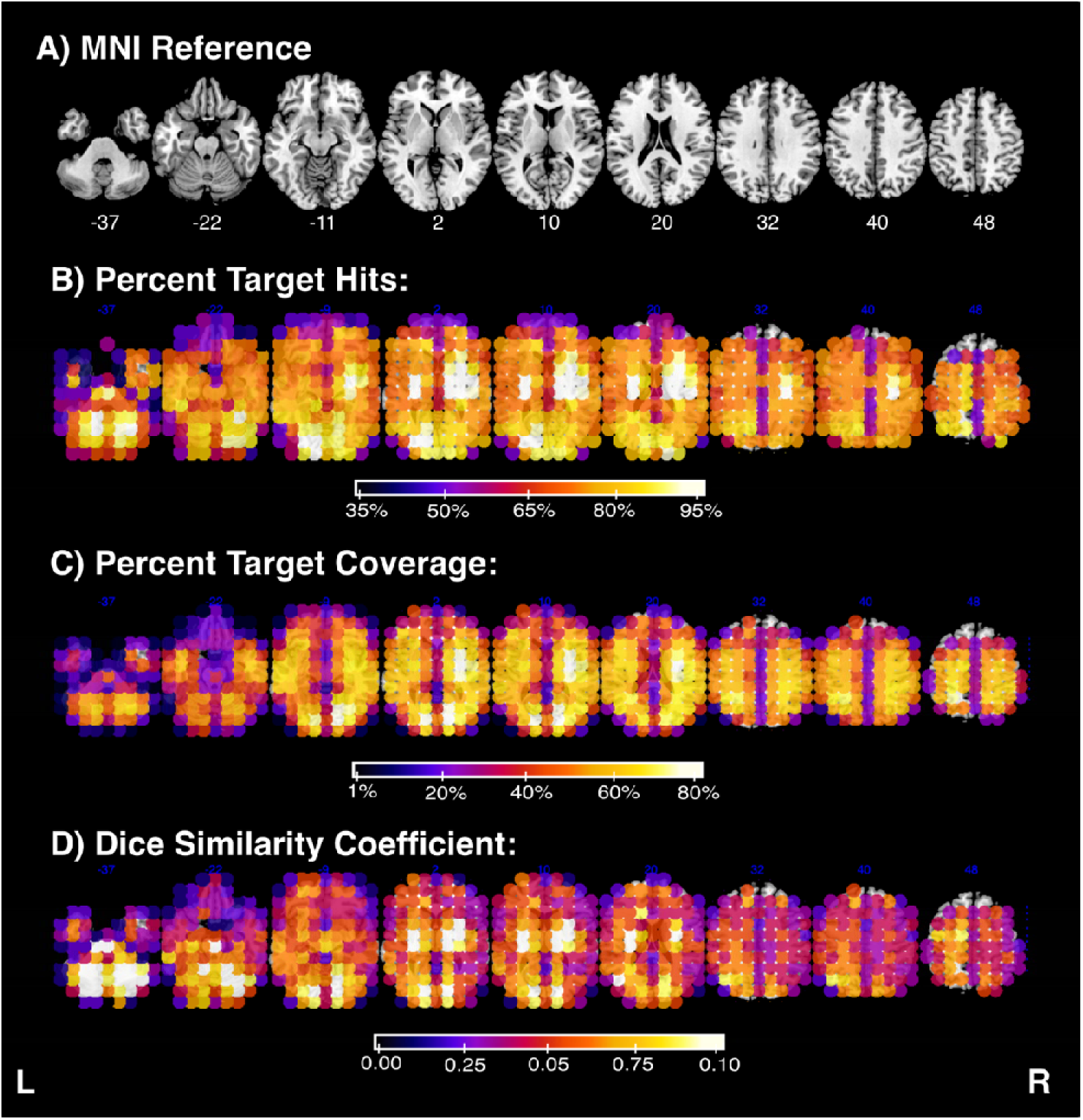
Lesion symptom mapping accuracy across different anatomical target areas. A) Provides a MNI reference to aid interpretation of subsequent panels. B) The percent of analyses which yielded significant results overlapping with the target for each considered anatomical target. C) The percent of target voxels which were identified as significant for each considered anatomical target. D) Dice similarity coefficient between significant voxels and underlying anatomical target voxels. Colour denotes mean accuracy for all analyses employing each defined anatomical target. Summary statistics reported at each target only include data from simulated analyses using the relevant target to simulate behaviour. L = left, R = right.

Across all univariate analyses yielding significant results, multivariate regression revealed that results accuracy (both in terms of coverage and Dice) was significantly modulated by sample size, average sample lesion volume, number of lesions impacting the target, and significant results cluster size (Dice Model: F(4,253204) = 3796, p < 0.001, R^2^ = 0.056, Percent Coverage Model: F(4,253204) = 03.39 × 10^4^, p < 0.001, R^2^ = 0.349) (see Figure 3).

**Figure 3:**
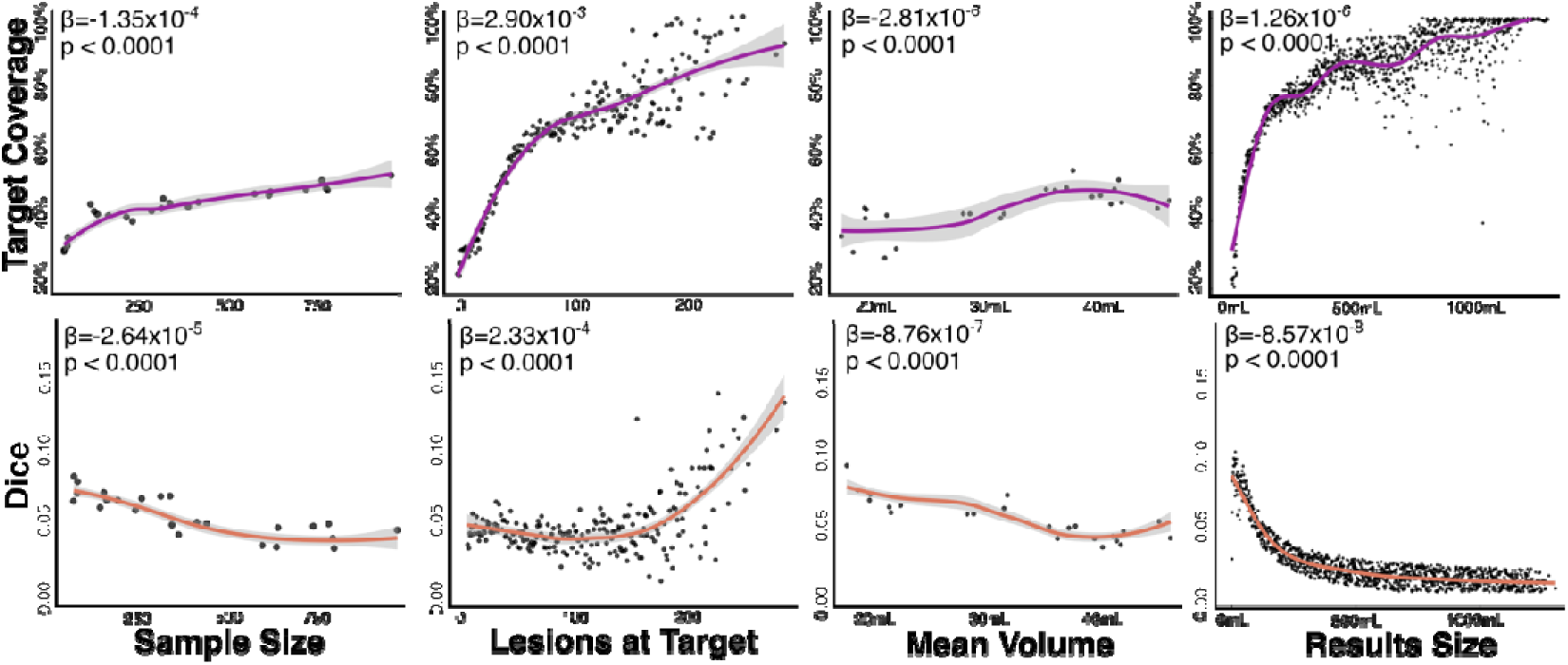
The relationship between key sample/results characteristics and analysis accuracy. Considered analysis factors are listed across the x-axis, while different accuracy measures (Coverage, Dice) are on the y-axis. The coefficients and significance for each factor within the described multivariate regression analyses are listed in the top left of each panel, while each panel depicts the best-fit general linear model for each individual factor.

### Impact of Individual Univariate Analysis Choices

Table 1 reports analysis accuracy descriptive statistics for each considered univariate analysis factor. Regression analyses outperformed chi-squared analyses in terms of both target coverage and Dice. Chi-squared analyses were more likely to yield significant results than regression analyses but were less likely to overlap with the target region. When considered alongside general analysis factors, the use of regression analyses positively predicted accuracy in terms of coverage (t = 48.13, p < 0.001, AIC change = 2304) and Dice (t = 69.04, p < 0.001, AIC change = 4721).

**Table 1:** Analysis accuracy across different analysis design factors. Value means and standard deviations (in parentheses) are presented for all univariate analyses employing each design factor. Sample Size reports the mean number of patients included in analyses and volume reports the average lesion volume within this sample. Per Sig reports the proportion of analyses which yielded significant results. Per Hit reports the proportion of significant results which overlapped with the underlying anatomical target. Dice and Coverage report the mean accuracy of analyses yielding significant results in each category. Analysis factors yielding the highest accuracy (in terms of dice and target coverage) are highlighted in red. P-values report results of ANOVA analyses comparing each value across all possible analysis design choices within each category.

|  | Sample Size | Volume | Per Sig | Per Hit | Results Size | Dice | Coverage |
| --- | --- | --- | --- | --- | --- | --- | --- |
| **Stroke Side** |  |  |  |  |  |  |  |
| Unselected | 612 (285) | 34.0 (8.1) | 69.30% | 73.04% | 140.5 (182.7) | 0.04 (0.07) | **0.471 (0.424)** |
| L | 235 (92) | 27.2 (4.9) | 68.11% | 70.99% | 54.3 (70.3) | **0.07 (0.09)** | 0.376 (0.396) |
| R | 323 (123) | 38.8 (8.2) | 58.71% | 71.55% | 91.7 (126.0) | 0.05 (0.08) | 0.403 (0.405) |
| Unilateral | 539 (237) | 32.7 (7.7) | 68.99% | 71.87% | 136.1 (178.9) | 0.04 (0.07) | 0.452 (0.420) |
| | $p < 0.001$ | $p < 0.001$ | $p < 0.001$ | $p < 0.001$ | $p < 0.001$ | $p < 0.001$ | $p < 0.001$ |
| **Scan Type** |  |  |  |  |  |  |  |
| Unselected | 625 (223) | 35.8 (4.0) | 71.73% | 72.50% | 133.9 (176.0) | 0.045 (0.071) | **0.457 (0.424)** |
| CT | 506 (179) | 39.6 (4.3) | 72.19% | 71.98% | 122.9 (163.8) | 0.046 (0.072) | 0.447 (0.422) |
| MR | 130 (43.2) | 20.2 (1.3) | 49.50% | 71.08% | 51.6 (77.8) | **0.071 (0.098)** | 0.359 (0.373) |
| | $p < 0.001$ | $p < 0.001$ | $p < 0.001$ | $p < 0.001$ | $p < 0.001$ | $p < 0.001$ | $p < 0.001$ |
| **Stroke Number** |  |  |  |  |  |  |  |
| Unselected | 490 (282) | 33.4 (8.3) | 66.73% | 72.55% | 116.4 (162.1) | **0.050 (0.078)** | **0.440 (0.417)** |
| First Time Only | 437 (237) | 33.7 (8.5) | 66.61% | 71.52% | 113.6 (159.0) | 0.059 (0.077) | 0.432 (0.415) |
| | $p < 0.001$ | $p < 0.001$ | $p < 0.001$ | $p < 0.001$ | $p < 0.001$ | $p = 0.083$ | $p < 0.001$ |
| **Analysis Type** |  |  |  |  |  |  |  |
| Chi Squared | 464 (262) | 33.6 (8.4) | 73.09% | 68.71% | 131.5 (144.6) | 0.028 (0.043) | 0.390 (0.407) |
| Regression | 464 (262) | 33.6 (8.4) | 65.25% | 72.87% | 110.9 (164.1) | **0.055 (0.083)** | **0.447 (0.417)** |
| | $p = 0.973$ | $p = 0.862$ | $p < 0.001$ | $p < 0.001$ | $p < 0.001$ | $p < 0.001$ | $p < 0.001$ |
| **Overlap Threshold** |  |  |  |  |  |  |  |
| >0 | 464 (262) | 33.6 (8.4) | 87.68% | 99.83% | 165.5 (188) | **0.083 (0.094)** | **0.698 (0.328)** |
| >10 | 464 (262) | 33.6 (8.4) | 71.11% | 66.67% | 110.9 (140.9) | 0.029 (0.048) | 0.334 (0.395) |
| >10% | 464 (262) | 33.6 (8.4) | 41.20% | 22.15% | 14.7 (21.1) | 0.014 (0.047) | 0.0525 (0.173) |
| | $p = 0.999$ | $p = 0.982$ | $p < 0.001$ | $p < 0.001$ | $p < 0.001$ | $p < 0.001$ | $p < 0.001$ |
| **Multiple Comparison Corrections** |  |  |  |  |  |  |  |
| Bonferroni | 464 (262) | 33.6 (8.4) | 68.97% | 72.65% | 89.5 (94.0) | 0.040 (0.051) | 0.446 (0.417) |
| FDR | 464 (262) | 33.6 (8.4) | 71.22% | 70.44% | 188.6 (214.9) | 0.024 (0.038) | **0.448 (0.424)** |
| Permutation | 464 (262) | 33.6 (8.4) | 57.55% | 73.70% | 34.6 (52.9) | **0.106 (0.121)** | 0.399 (0.397) |
| | $p = 0.993$ | $p = 0.837$ | $p < 0.001$ | $p < 0.001$ | $p < 0.001$ | $p < 0.001$ | $p < 0.001$ |
| **Lesion Volume Corrections** |  |  |  |  |  |  |  |
| Regression | 464 (262) | 33.6 (8.4) | 68.04% | 70.80% | 129.0 (217.6) | 0.055 (0.086) | 0.435 (0.419) |
| None | 464 (262) | 33.6 (8.4) | 70.19% | 68.0% | 118.4 (144.5) | 0.042 (0.073) | 0.421 (0.421) |
| DTLVC | 464 (262) | 33.6 (8.4) | 57.50% | 81.21% | 80.4 (92.7) | **0.071 (0.089)** | **0.493 (0.407)** |
| | $p = 1.00$ | $p = 0.997$ | $p < 0.001$ | $p < 0.001$ | $p < 0.001$ | $p < 0.001$ | $p < 0.001$ |

### Impact of Sample Inclusion Criteria Choices

Samples including all lesions (regardless of location) had the highest target coverage while samples including only left hemisphere patients yielded the highest Dice coefficient (Table 1). When stroke side categories were added to the regression models predicting accuracy from general factors (described above), employing all lesions (regardless of location) significantly improved target coverage (t = 10.52, p < 0.001, AIC change = 109) and Dice (t = 5.96, p < 0.001, AIC change = 34). No other lesion location criteria significantly improved target coverage or Dice.

Analyses including all patients, regardless of imaging modality, yielded the highest target coverage, while analyses using only MR resulted in the highest Dice. When scan type inclusion criteria were incorporated into regression models predicting accuracy, using only CT scans improved analysis accuracy both in terms of target coverage (t = 9.914, p < 0.001, AIC change = 96) and Dice (t = 4.733, p < 0.001, AIC change = 21). Notably, MR-only analyses had smaller sample sizes and were significantly more likely to produce false negative results (see Table 1). However, in cases where MR analyses did yield significant results they produced comparatively high Dice.

Including all lesions (regardless of lesion numerosity) yielded higher target coverage than first-time stroke samples, but Dice was not different between these groups. Including the number of strokes for each individual as a covariate did not significantly add to models predicting target coverage from general analysis factors (Coverage: t = 0.374, p = 0.708, AIC change = 2). However, using all lesion samples (rather than first stroke only) was associated with improved Dice (t = 3.99, p < 0.001, AIC change = 14).

Overall, using all lesions (regardless of location), employing only CT-derived lesions, and including all patients (regardless of stroke numerosity), were the best-performing analysis choices.

### Impact of Analysis Parameter Choices

Analyses which did not employ a minimum overlap threshold significantly outperformed those employing a minimum overlap threshold in terms of both Dice and target overlap. This finding cannot be a secondary consequence of increased results cluster sizes, as the use of no minimum overlap threshold positively correlated with both Dice (t = 378.36, p < 0.001, AIC change= 64,187) and target coverage (t = 270.29, p < 0.001, AIC change = 113,469) when general analysis factors were controlled for. No other overlap threshold criteria significantly improved target coverage or Dice.

FWER permutation corrections yielded the highest Dice, while FDR corrections produced the best target coverage. Notably, analyses using FDR corrections produced clusters which were much larger than those produced by analyses using FWER permutation or Bonferroni corrections. When general analysis factors (including the size of the results cluster) were controlled for, the use of FDR corrections predicted worse target coverage (t = −103.77, p < 0.001, AIC change = 10,543) and worse Dice (t = −114.33, p < 0.001, AIC change = 12,743). Conversely, the use of FWER permutations predicted improved coverage (t = 52.18, p < 0.001, AIC change = 2,706) and Dice (t = 209.92, p < 0.001, AIC change = 40,625) when general analysis factors were considered. The use of Bonferroni corrections predicted better target coverage (t = 51.24, p <0.001, AIC change = 2611), but lower Dice (t = −59.55, p < 0.001) when general analysis factors were controlled for. For this reason, FWER permutation corrections were selected as the best-performing multiple comparison correction option.

The use of DTLVC lesion volume corrections yielded the best analysis accuracy in terms of both Dice and target coverage. The use of DLTVC corrections predicted improved analysis accuracy (over and above general analysis factors) in terms of both target coverage (t = 94.22, p < 0.001, AIC change = 1045) and Dice (t = 48.77, p < 0.001, AIC change = 2655). Regressing out lesion volume predicted improved Dice (t = 5.79, p < 0.001, AIC change = 32), but did not improve target coverage. DLTVC corrections were therefore selected as the best-performing lesion volume correction.

Within analysis parameter factors, employing DTLVC lesion volume corrections, using FWER permutation corrections for multiple comparisons, and not using minimum lesion overlap analysis inclusion thresholds were found to yeild the most accurate results.

### Best-Performing Models

Univariate analyses employing the best-performing combination of analysis type, sample inclusion criteria, and analysis parameters (described above) were evaluated relative to analyses using other factor combinations. Of the 667 analyses that were run using the best-performing analysis design, 83.5% yielded significant results 99.6% of which overlapped with the target. These significant analyses yielded an average Dice of 0.21 (SD = 0.12, range = 0 – 0.59) and an average coverage of 68.4% (SD = 31.9%, range = 0 – 100%). Each of these scores was significantly higher within the best-performing model relative to all other analysis designs (percent significant: X2=44.4, p < 001; percent hits: X2 = 263.4, p < 0.001; Dice: t = 27.2, p < 0.001; Coverage: t = 17.3, p<0.001). Notably, mean dice was 4.25 times higher and coverage was 1.57 times higher within significant models using the best combination of factors relative to all other models.

### Multivariate analysis simulation results

60.7% (n = 3665) of the potential multivariate analysis designs met criteria for inclusion in the simulation analysis. These accuracy data were compared with those of the univariate analyses using identical sample inclusion criteria, analysis parameters, and target ROIs. Multivariate analyses yielded a lower proportion of significant results relative to matched univariate analyses (63.0% vs 79.8%, X^2^ = 2916.4, p <0.001), and these results were less likely to overlap with the underlying target (44.2% target hits versus 52.7%, X^2^ = 52.5, p <0.001). Multivariate analyses yielded higher Dice relative to univariate analyses (0.068 vs. 0.025, t(2459.4) = 16.82, p < 0.001), but produced lower target overlap (0.15 vs. 0.24, t(4652.1) = −13.057, p < 0.001). These relationships remained consistent across different considered analysis designs (Table 2 & 3). Accuracy differences between SCCAN and univariate approaches varied across different target ROIs (Figure 4), with the largest improvements in Dice coefficient in multivariate relative to univariate analyses present in areas with the most lesion coverage.

**Figure 4:**
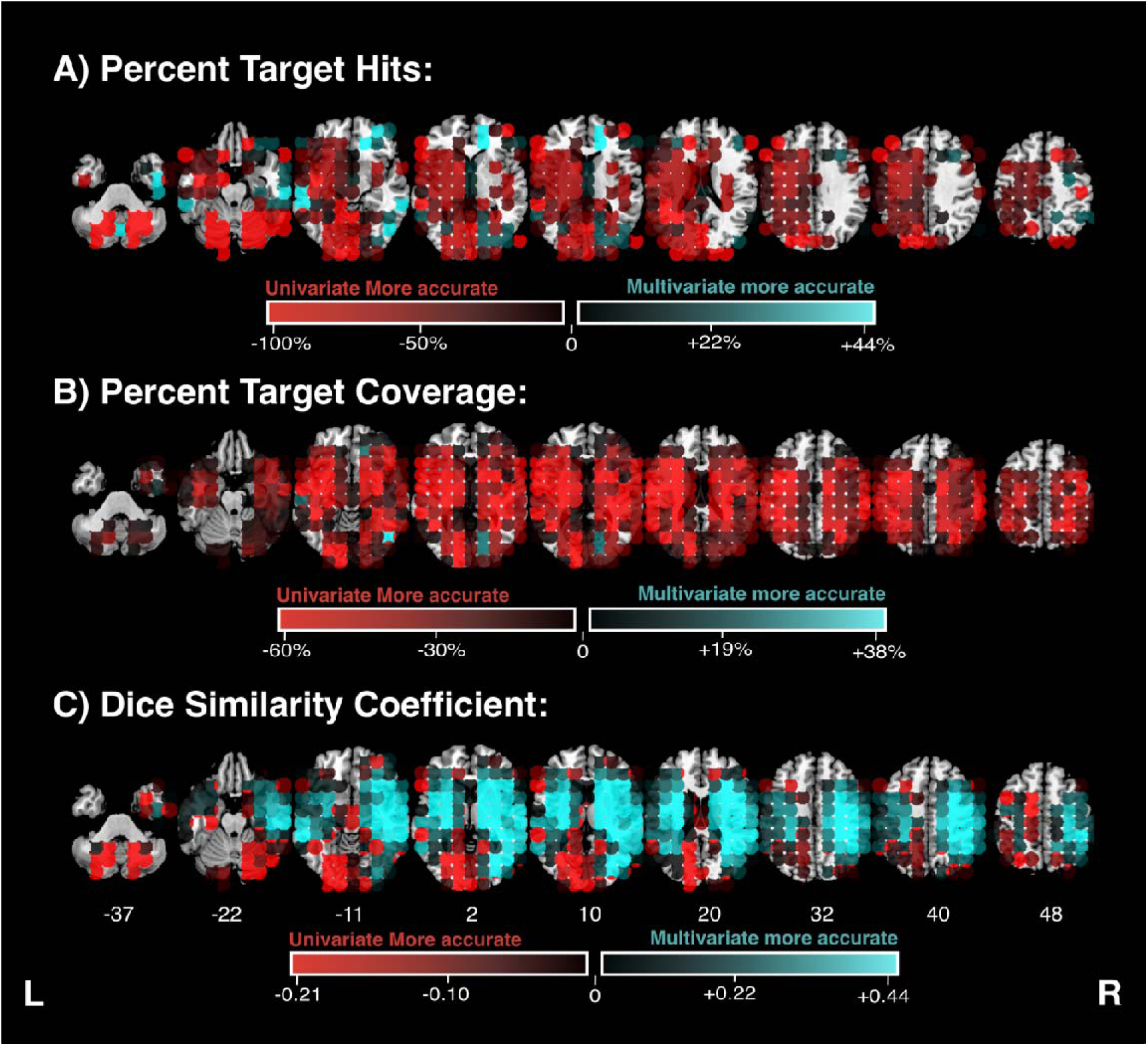
A comparison of lesion symptom mapping accuracy between multivariate (SCCAN) and univariate analysis approaches. Lesion symptom mapping accuracy across different anatomical target areas. Colour denotes mean accuracy for all analyses employing each defined anatomical target. Accuracy is quantified in terms of percent of analyses which yielded significant results overlapping with the target (Panel B), the percent of target voxels which were identified as significant (Panel C), and dice similarity coefficient between significant voxels and the underlying anatomical target (Panel D). L = left, R = right.

**Table 2:** SCCAN versus regression (univariate) lesion symptom mapping accuracy across different analysis factors. Value means (for SCCAN analyses) are presented alongside the difference between SCCAN and univariate regression performance (in parentheses). Positive values represent cases in which values in SCCAN analyses were higher relative to univariate analyses and negative values represent cases in which SCCAN values were lower. Per Sig reports the proportion of analyses which yielded significant results. Per Hit reports the proportion of significant results which overlapped with the underlying anatomical target. Dice and Coverage report the mean accuracy of analyses yielding significant results in each category.

| Stroke Side |  | Per Sig | Per Hit | Results Size | Dice | Coverage |
| --- | --- | --- | --- | --- | --- | --- |
|  | L | 60.3% (-13.9) | 46% (-4.3) | 15.0 (-25.7) | 0.069 (+0.035) | 0.146 (-0.062) |
|  | R | 64.9% (-17.4) | 43.1% (-12.8) | 25.7 (-46.5) | 0.067 (+0.042) | 0.153 (-0.115) |
| Overlap Threshold |  |  |  |  |  |  |
|  | >10 | 70.2% (-26.0) | 61.7% (-6.6) | 35.7 (-43.3) | 0.088 (+0.053) | 0.228 (-0.107) |
|  | >10% | 55.8% (-4.7) | 22.5% (-6.8) | 3.6 (-20.3) | 0.044 (+0.028) | 0.053 (-0.035) |
| Lesion Volume Corrections |  |  |  |  |  |  |
|  | Regression | 63.8% (-11.6) | 39.8% (-15.6) | 33.9 (-10.2) | 0.071 (+0.035) | 0.158 (-0.091) |
|  | None | 62.1% (-19.4) | 48.8% (-2.3) | 8.71 (-61.7) | 0.065 (+0.045) | 0.142 (-0.090) |

**Table 3:** SCCAN versus chi-squared (univariate) lesion symptom mapping accuracy across different analysis factors. Value means (for SCCAN analyses) are presented alongside the difference between SCCAN and univariate chi-squared performance (in parentheses). Positive values represent cases in which values in SCCAN analyses were higher relative to univariate analyses and negative values represent cases in which SCCAN values were lower. Per Sig reports the proportion of analyses which yielded significant results. Per Hit reports the proportion of significant results which overlapped with the underlying anatomical target. Dice and Coverage report the mean accuracy of analyses yielding significant results in each category.

| Stroke Side |  | Per Sig | Per Hit | Results Size | Dice | Coverage |
| --- | --- | --- | --- | --- | --- | --- |
|  | L | 60.3% (-17.5) | 46% (-2.2) | 15.0 (-36.9) | 0.069 (+0.047) | 0.146 (-0.053) |
|  | R | 64.9% (-23.9) | 43.1% (-9.6) | 25.7 (-78.2) | 0.067 (+0.053) | 0.153 (-0.099) |
| Overlap Threshold |  |  |  |  |  |  |
|  | >10 | 70.2% (-28.6) | 61.7% (-6.4) | 35.7 (-77.2) | 0.088 (+0.065) | 0.228 (-0.116) |
|  | >10% | 55.8% (-13.0) | 22.5% (-3.2) | 3.6 (-30.8) | 0.044 (+0.033) | 0.053 (-0.024) |

## Discussion

We have described a comprehensive investigation into how analysis design factors impact the accuracy of lesion symptom mapping analyses. This large-scale simulation design provides novel insight into how a wide range of plausible analysis designs, samples, and underlying anatomical targets may interact to modulate the sensitivity and specificity of lesion symptom mapping analyses. Across all simulated designs, lesion symptom mapping analysis accuracy was highly variable and substantially modulated by the specific parameters employed in each analysis. These results provide important practical guidance by highlighting differential sample inclusion, analysis parameter, and analysis type choices which can be employed by authors aiming to maximise the specificity or sensitivity of lesion mapping analyses. Taken together, the findings demonstrate that there is no single optimal study design but that each individual analysis choice has advantages and disadvantages which must be considered in the context of a study’s individual aims.

### Factors which reliably improve or reduce accuracy in lesion symptom mapping

Despite the wide variability in accuracy amongst individual design choices, the results highlight several key factors which were consistently associated with either reduced or improved accuracy. First, analyses with better lesion coverage generated better accuracy. In univariate analyses, percent coverage of target ROIs improved dramatically in samples with >50 lesions at the target relative to analyses with fewer “impaired” patients. This effect was less pronounced when accuracy was measured in terms of Dice, but a clear improvement was notable in analyses that included >150 “impaired” patients. This finding builds upon past work that established the importance of large sample sizes in lesion symptom mapping studies (Lorca-Puls et al., 2018; Moore, Demeyere, et al., 2023). From first principles, statistical power is optimal where regions are impacted in half of the included sample. In real-world lesion symptom mapping analyses, the degree of lesion overlap at the target correlate is a function of the overall sample size and the number of patients within the sample who are impaired on the behavioural measure of interest. We found a negligible impact of including patients based on multiple stroke lesions and a clear benefit of combining different neuroimaging modalities in lesion symptom mapping. These results suggest that future studies could employ more lenient inclusion criteria to maximise the number of patients included in analyses.

For the univariate analyses, approaches using direct total lesion volume controls (DLVTC) reliably outperformed other volume correction options in terms of both Dice and target coverage. In line with past work, regression-based volume corrections were more accurate than analyses with no volume corrections, but yielded less accurate results relative to DLVTC approaches (DeMarco & Turkeltaub, 2018; Zhang et al., 2014). Past work has theorised that regression-based corrections may be too conservative for lesion symptom mapping because critical variability relating to true correlates may be regressed out alongside lesion volume (Sperber, 2022). The results of our study align with this hypothesis, as DLVTC approaches are less conservative and were found to generate more accurate results. Notably, many popular lesion symptom mapping software toolkits do not offer DLVTC as a built-in correction method. Given the demonstrated benefit of DLVTC approaches, researchers should aim to use toolkits which offer the option to use this approach, and developers should aim to integrate this option into new and existing lesion symptom mapping analysis packages.

Employing a minimum voxel-level inclusion criterion of 10% of the total sample was associated with reduced performance across all considered designs and accuracy measures. Studies using 10% overlap inclusion thresholds were less likely to generate significant results and, in cases where significant results were produced, were less accurate than the other tested cutoffs. Employing minimum overlap thresholds is intended to reduce spatial displacement of lesion symptom mapping results (Karnath et al., 2018; Sperber & Karnath, 2017). Spatial bias is generally strongest within cortical regions, meaning that the inclusion of many subcortical targets in our analysis may have masked any potential accuracy benefit from employing stricter minimum overlap thresholds (Karnath et al., 2018; Sperber & Karnath, 2017). The use of 10% overlap inclusion thresholds is also more commonly used in studies with smaller samples (<50) relative to larger-scale investigations which frequently opt for minimum inclusion cut-offs which do not scale with sample size (e.g., >3, >5, >10). It is plausible that using a 10% overlap inclusion threshold is more poorly suited to lesion mapping studies with larger samples (>100) relative to smaller samples (<50), as the number of overlapping parcels needed to meet a percentage-based cut-off will increase as sample size increases. This possibility was not explicitly evaluated in our study as all included samples contained at least 67 patients. Where possible, lesion symptom mapping analyses should ideally include a minimum of 100 patients (Moore et al., 2023). In this context, the results of the current study suggest that a 10% overlap inclusion criterion may not be optimal in lesion symptom mapping studies.

Within univariate analyses, chi-squared approaches were consistently less accurate than regression-based approaches. Chi-squared approaches were more likely to generate significant results, but these results were less likely to contain the target region and were less accurate. We binarised simulated behavioural scores for chi-squared analyses using a single threshold (0.50). It is unclear whether the results reported here would remain consistent across all potential impairment thresholds, but it seems plausible that approaches that constrain variability in input behavioural scores may be limited in their ability to precisely localise brain-behaviour relationships (de Haan & Karnath, 2018). There are also several alternative statistical approaches to handle binarised behavioural data, such as analyses using Liebermeister measures as the base statistical test (Rorden et al., 2007). Given that behavioural data from patient samples is often non-parametric, it is important to evaluate what the optimal method for handling categorical and/or binary scores in lesion symptom mapping may be.

### Key Differences from previous lesion symptom mapping work

This results of our study are largely in line with previous work, but there are several notable exceptions. First, analyses that did not employ a minimum overlap threshold generated more accurate results in terms of percent hits, Dice, and target coverage. This result is surprising as previous simulation work has concluded that applying a minimum overlap inclusion threshold improves lesion symptom mapping accuracy (Sperber & Karnath, 2017). It is plausible that minimum overlap thresholds may differentially interact with other correction factors. For example, significance thresholds for analyses using some correction approaches (e.g. Bonferroni) may become stricter when minimum overlap thresholds are not used, as this would increase the number of tested voxels.

Analyses which employed only lesions derived from CT imaging consistently outperformed analyses which used a combination of MR and CT or MR only. This result is surprising for two reasons: first, because selecting based on imaging type reduces sample size (see Table 1), and second because MR is generally more sensitive to lesion damage (Brazzelli et al., 2009; Kloska et al., 2004). Within MR, specific modalities such as T2 and FLAIR are more well suited to lesion segmentation than others (e.g. T1) (de Haan & Karnath, 2018; Moore et al., 2023). Most of the data used in the current study came from UK stroke settings where MR imaging is generally only used in special cases where patients have small lesions which are difficult to localise using CT or are neurologically complex cases (e.g., comorbid neurological abnormalities) (Lövblad et al., 2015; Wardlaw et al., 2004; Wardlaw, Seymour, et al., 2004). This sample difference is reflected in the finding that the MR imaging used in this study depicts smaller lesions with a qualitatively different spatial topography than CT imaging (see Group 20 vs Group 22 in Supplementary Figure 2). It is therefore plausible this distribution difference may have negatively impacted analysis accuracy, but it is unclear whether this effect would be expected to generalise to other simulated or real-world lesion analyses.

### Univariate versus multivariate lesion symptom mapping analyses

Multivariate analyses consistently outperformed univariate analyses in terms of Dice, but univariate analyses were reliably more accurate in terms of target coverage. This result is in line with past work indicating that there is no universal superiority of one analysis type over another, but that each analysis type offers specific advantages and weaknesses which must be considered in the context of individual study goals (Brazzelli et al., 2009).

Past work which has reported that multivariate analyses consistently outperform univariate analyses has generally quantified accuracy in terms of measures such as Dice coefficient and results displacement distance (Mah et al., 2014; Pustina, 2019). Our study demonstrates that these measures may not fully capture the performance of lesion symptom mapping analyses, because improvements in Dice scores were generally associated with decreases in target coverage. Results displacement is strongly associated with the size of significant voxel clusters (Moore et al., 2023), meaning that displacement may be comparatively higher in univariate versus multivariate analyses solely due to the comparatively larger significant voxel clusters generated by univariate analyses.

Additionally, lesion symptom mapping accuracy was often low even within the best-performing univariate and multivariate analysis designs. This result indicates that neither multivariate nor univariate lesion symptom mapping analysis designs should be assumed to be perfectly accurate, as both analysis designs are susceptible to voxel-wise false positives and misses due to factors such as variable lesion overlay (e.g. statistical power) and spatial biases in lesion distributions (Kimberg et al., 2007; Mah et al., 2014; Moore, Demeyere, et al., 2023). Overall, these findings are in line with previous work reporting that multivariate and univariate approaches both have benefits, and that the optimal analysis type depends on the specific goals of individual analyses.

### Sensitivity versus Specificity in Lesion Symptom Mapping

Lesion symptom mapping, like any statistical approach, entails an inherent trade-off between sensitivity and specificity. The results of the current study provide clear guidance to authors aiming to design lesion symptom mapping analyses which prioritise sensitivity or specificity while also highlighting the trade-offs associated with each approach. Studies which aim to maximise sensitivity prioritise detecting correlates (if they exist) over minimising voxel-wise false positives. In this case, it is methodologically valid for authors to adopt more liberal correction approaches that maximise the probability of generating results which overlap with the underlying target (e.g., univariate analysis designs, FDR corrections, no corrections for lesion volume). However, this approach limits the degree to which the identified anatomy can be assumed to be functionally related to the deficit of interest due to its high voxel-wise false positive rate. Studies prioritising sensitivity should therefore strictly limit theoretical interpretation and causal inferences pertaining to identified correlates, and clearly acknowledge the elevated risk of voxel-wise false positives. Sensitivity-driven analyses can instead focus on describing broad lesion patterns associated with the behaviour of interest and qualitatively comparing these patterns across patient groups. This approach may be most appropriate in cases where samples are small, and where impairments are rare and descriptive.

Conversely, studies which aim to prioritise specificity may adopt more conservative statistical approaches (e.g. multivariate analyses, permutation-corrections, DLVTC lesion volume corrections). Specificity-driven studies can be more confident that identified regions are causally involved in the underlying function of interest, while acknowledging that some critical regions may be missed. This approach is ideal for studies aiming to make causal interpretations about identified neural correlates, because significant results arising from this more rigorous analytical approach provide stronger evidence in support of causal brain-behaviour relationships than more liberal statistical approaches.

Sensitivity-driven and specificity-driven analysis designs are not mutually exclusive; indeed, it is feasible to use both analytical approaches in tandem. Correlates which remain significant in both sensitivity-driven and specificity-driven analytical designs can be confidently interpreted as being causally involved with the behaviour of interest (Ivanova et al., 2021). In cases where significant results do not emerge from strict, specificity-driven analyses, the results of sensitivity-driven analyses can be used to identify the presence of broader lesion patterns which may distinguish between behavioural groups of interest. This combined analytical approach is in line with past work suggesting that combining both multivariate and univariate lesion symptom mapping analyses within individual studies may be the most informative way to identify and understand the role of implicated neural correlates (Ivanova et al., 2021). The present study adds to this previous work by identifying a range of additional parameters, outside of univariate/multivariate approaches, which can be used to tailor analyses toward prioritising voxel-wise sensitivity and specificity.

### Limitations

There are several factors that may modulate lesion symptom mapping accuracy which were not evaluated in this study. These include several common statistical tests (e.g. Liebermiester tests, t-tests, support vector regression) as well as important factors (e.g. sample size) which were not explicitly manipulated in this study. Additionally, our design prioritised voxel-level lesion symptom mapping approaches. It remains unclear whether the documented effects would also hold for disconnection-based approaches such as network- and tract-level lesion symptom mapping. Our study simulated behavioural deficits based on the degree to which real lesions overlap with single, spatially continuous brain ROIs. It is not possible to know whether the documented effects would remain consistent in cases where behaviour is supported by multiple, spatially distant regions or the connections between these regions. The target ROIs we used here are also not naturalistic as they were represented by voxel spheres which would not be expected to align with real-world functional ROI boundaries, or the overlap topographies present in lesion overlays. Future studies should aim to evaluate whether consistent results are found when more realistic potential underlying neural correlates are considered (e.g. functional ROIs). Previous work has demonstrated that adding noise to simulated behavioural scores can dramatically reduce the accuracy of lesion symptom mapping analyses (Ivanova et al., 2021). We did not include this manipulation, and therefore cannot evaluate whether different lesion symptom mapping analysis types are more susceptible to the presence of behavioural noise.

### Conclusions

We have presented a comprehensive investigation into how individual analysis choices modulate results accuracy in lesion symptom mapping studies. We identified several key analytical choices which were universally associated with improved lesion symptom mapping accuracy. However, the results of our study clearly illustrate that the optimal analysis design is ultimately dependent on the specific goals of individual studies. For this reason, it is important that lesion symptom mapping analysis parameters are explicitly selected to prioritise results sensitivity or specificity, and for any resultant significant voxel clusters to be interpreted with this design’s strengths and limitations in mind. We believe that this practice, when applied systematically, has the potential to improve both the quality and accuracy of lesion symptom mapping analyses as well as improving the reliability of theoretical inferences made based on lesion mapping results.

## Acknowledgements

We would like to thank the participants who took part in the Oxford Cognitive Screening study and all research staff who contributed towards data collection. In particular, we acknowledge the contributions to data collection made by Ms Ellie Slavkova, Ms Grace Chiu, and Ms Romina Basting.

## Author Credit Statement

In line with the CReDiT taxonomy, MJM was responsible for conceptualisation, software, data curation, formal analysis, methodology, visualisation, and writing (original draft). CR was responsible for writing (review & editing). GR was responsible for resources and writing (review & editing). JBM was responsible for supervision and writing (review & editing). ND was responsible for supervision, resources, and writing (review & editing).

