## Supplemental Materials for "Methodological choices strongly modulate the sensitivity and specificity of lesion-symptom mapping analyses"

**Supplementary:**

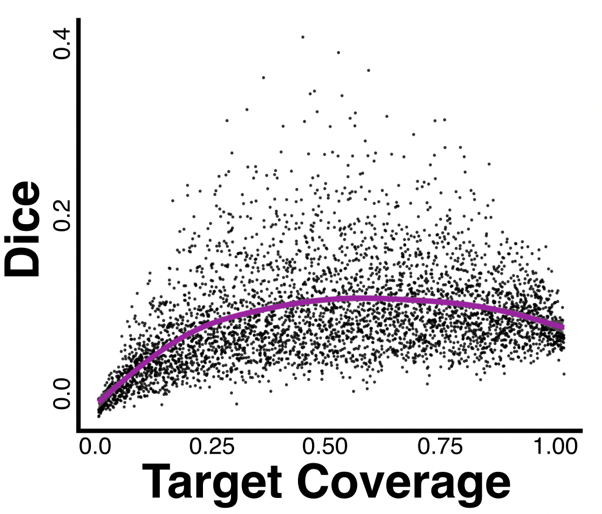

***Supplementary Figure 1:*** *The relationship between Dice coefficients and target percent coverage across all simulated univariate analyses which yielded significant results.*

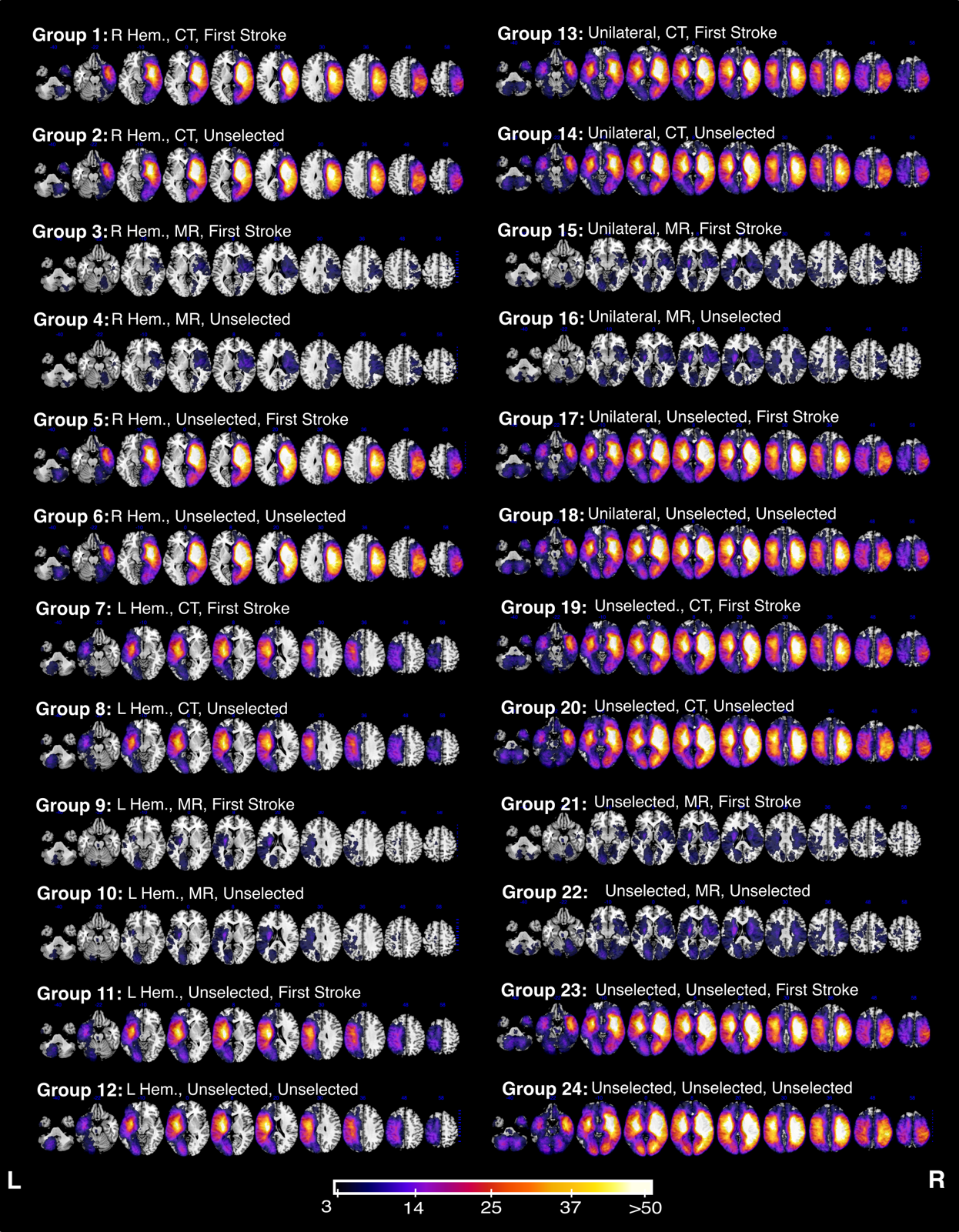

***Supplementary Figure 2:*** *Lesion overlays for each of the possible 24 patient groups. Colour denotes the number of lesions impacting each area. MNI axial slices between X = -40 – 58 are shown.*
